# Enhanced cell wall mechanics in VirtualLeaf enable realistic simulations of plant tissue dynamics

**DOI:** 10.1101/2024.08.01.605200

**Authors:** Richard W. van Nieuwenhoven, Bruno Hay Mele, Roeland M.H. Merks, Ruth Großeholz

**Author notes:** These authors contributed equally to this work (Richard W. van Nieuwenhoven, Ruth Großeholz).

## Abstract

Computational modelling has become essential to advancing our understanding of plant developmental and physiological processes, necessitating the development of new computational approaches and software. Here, we present VirtualLeaf-2.0, an updated version of this modelling framework for the biophysical and biomechanical interactions between cells in plant tissues, with novel features for more detailed modelling of the cell wall. In particular, the updated version of VirtualLeaf enables detailed modelling of variations in cell wall stability and cell wall sliding up to the level of individual cell wall elements. The plant cell wall plays a pivotal role in plant development and survival, with younger cells generally having thinner, more flexible (primary) walls than older cells. Cell wall stability is further affected by signalling in growth processes and pathogen infection. The improvements of VirtualLeaf lay the groundwork for using VirtualLeaf to address novel questions involving plant tissue dynamics during growth, tissue formation and pathogen defence, as illustrated with example simulations.

## 1 Introduction

Computational modelling has become an indispensable instrument in plant biological research. It has advanced the field from a descriptive approach of genotype-phenotype relations to an explanatory field where consequences of gene activity in the context of the tissue have been integrated, facilitating the understanding of these complex gene-to-phenotype systems [1, 2]. Computational models allow the translation of conceptual models of integrative, biological hypotheses into mathematical equations to generate and test different scenarios [3], and to identify crucial interactions [4]. Thus, combining molecular, mechanical and environmental factors into an integrated mathematical model generates a deeper understanding of plant growth and adaptation [5, 6, 7, 8, 9].

Several recent advances in plant research result from interdisciplinary studies that combine an iterative cycle of experiments and computational simulations. For example, computational models have predicted differential sensitivity to auxin and the buffering capacity of its signalling pathways [10]. A detailed kinetic model of the rapid brassinosteroid response demonstrated that the canonical signalling pathway requires the involvement of a cation channel to function [11]. Similarly, modelling has provided mechanistic hypotheses for the regulation and maintenance of the shoot apical meristem [12]. However, understanding how a gene regulatory network gives rise to a specific phenotype requires more than analysing the molecular network alone. Here, the broader context needs to be considered, such as cellular interactions, tissue layout, and a cell’s position within the tissue, which again determines the signals it receives. The need for the context of the whole tissue and organism was nicely demonstrated by Besnard *et al*. who analysed the complex feedback mechanisms involved in auxin-mediated organogenesis [13]. Models that integrate tissue layout and gene regulatory networks for auxin signalling have yielded insights into patterning during root and shoot development [10, 14, 15, 16, 17]. Mironova *et al*. showed that the regulation of PIN expression by auxin is sufficient to cause the auxin pattern in the root [14]. Vernoux *et al*. analysed the interplay of auxin response factors (ARF) and Aux/IAAs in shoot development for robust shoot patterning [10]. Given the complexity of living systems, these topics highlight that modelling is not merely a supplementary instrument but a fundamental approach to understanding the regulatory mechanisms governing plant development and survival [18].

Addressing these modelling questions, especially concerning the spatial context, requires a computational framework capable of analysing cellular signalling within the spatial context of the tissue. Multiple frameworks are available for animal tissues, including cellular Potts modelling frameworks Morpheus [19], CompuCell3D [20], and Tissue Simulation Toolkit [21], as well as off-the-lattice frameworks such as PhysiCell [22] and vertex-based models such as those implemented in TissueForge [23] and in CHASTE [24].

Modeling frameworks aimed at animal tissues have been used for plant tissues [25], but they do not properly represent the rigid cell wall, which prevents relative movement between cells under physiological conditions, the roles of turgor pressure in providing plant rigidity, and the role of water fluxes between cells [9, 26]. Therefore, several tools specific to plant tissue simulations have been developed in recent years. MorphoGraphX provides a framework in which the mechanical aspects of tissue formation can be analysed in 3D [27], e.g. to investigate the shape of puzzle cells [8]. The second generation MorphoDynamX [28] and its extension MorphoMechanX [29] also integrate the simulation of cellular dynamics and Finite Element Method (FEM) for solid mechanics [30, 31]. CellModeller provides an open-source 2D framework for simulating cellular tissue layouts and diffusion of morphogens [32]. Other modelling approaches concerning plant tissues provide a framework for the 3D mechanical analysis of plant tissues at single cell resolution [33]. A more recent study linked water fluxes with cell wall mechanics during tissue growth computationally [9]. In the present work, we will be extending VirtualLeaf [34], a modelling framework that is, like some of the frameworks above, rooted in the idea of cell-based modelling (CBM).

The key idea of CBM is to predict how behaviours of single cells and their response to chemical or mechanical signals from the micro-environment (the input to a CBM), gives rise to the behaviour and pattering of the whole tissue, organ, or organism, and, importantly, how the resulting patterns feed back on the behaviour of the single cells [35, 36]. CBM describes individual cells as spatial entities with defined mechanical properties that interact according to pre-defined rules and thereby enables the analysis of feedback and communication mechanisms across different biological scales. This approach enables the analysis of dynamic changes in cell shape, division patterns, and cell polarity. CBM can thus capture the influence of turgor pressure and differential growth rates. By combining cellular interactions with intra- and intercellular signalling processes, CBM, using VirtualLeaf or its derivative codes, has provided insights into pattern formation [5], vascular tissue differentiation [6], and growth regulation [7].

While all of these frameworks have their special area of applicability and their own strengths and weaknesses, there is still a crucial step missing in order to accurately model plant tissue growth and development. When analysing plant tissues using computational modelling, it is crucial to consider the specific characteristics of plant morphology: Plant tissue development and integrity rely in part on the precise regulation of cell wall stability and properties [37, 38]. On the one hand, the cell wall needs to be more flexible during growth to allow cellular growth [39]. Excessive cell wall weakening can compromise plant integrity and render the plant susceptible to pathogen infections [40]. Cell wall properties generally vary significantly between cell types and between older and younger cells — younger cells have thin, more flexible (primary) cell walls [41], and older cells generally acquire thicker, more stable (secondary) cell walls over time [42]. The composition of the cell wall is influenced by multiple factors, such as the age of the cell, its biophysical properties, and anisotropies [43, 44]. Various signalling pathways affect and regulate cell wall stability, e.g. auxin and brassinosteroids during growth [45, 46, 47, 48, 49]. Plant physiological processes can likewise increase the stability of the cell wall, e.g. in response to elevated mechanical stress [50] and in defence against pathogens [51]. as the cell wall serves as a mechanical barrier that pathogens must overcome to become transmissible between cells [52, 53]. Under particular circumstances, plants actively prompt cell wall remodelling, a complex process that involves the rearrangement of cells in a specific tissue, e.g. during lateral root emergence [54].

In an effort to provide a framework for more realistic plant tissue simulations, where the biomechanical consequences of signalling processes can be mapped accurately, we now present an updated version of VirtualLeaf. VirtualLeaf is a CBM-based system where intra- and intercellular signalling processes can be simulated and analysed in the context of the tissue layout in 2D, considering cellular interactions, cell differentiation and cell growth. VirtualLeaf simulations have already yielded valuable insight into auxin patterning [55, 56, 57], leaf development [58], vascular patterning [6, 59], root elongation[60]. However, a more detailed representation of mechanical interactions is required to achieve realistic simulations of plant tissues. Current VirtualLeaf models assume that plant cells are rigidly connected. This updated version of VirtualLeaf introduces limited cellular migration along neighbouring cells under specific conditions, allowing the simulation of developmental processes that involve cell wall remodelling, such as lateral root emergence. We further implemented realistic cell wall mechanics, taking into account the plastic-elastic properties of cell wall filaments. We further implemented limited cellular migration along neighbouring cells under specific biochemical conditions that support the simulation of cell wall remodelling events such as lateral root emergence. Together, these additions to VirtualLeaf provide a crucial step forward in accurately simulating plant tissue development. In the future, VirtualLeaf will need to support 3D simulations as well, in line with other widely used tools such as MorphoDynamX and MorphoGraphX, as 3D simulations are necessary for a comprehensive understanding of plant developmental processes. In the meantime, this new version of VirtualLeaf represents an important advancement by addressing the need of the community for a realistic framework that incorporates physiologically accurate mechanics and cell wall remodelling in a 2D context. The 2D version further provides a way to screen parameters while avoiding significant computational costs [6].

Furthermore, the newly introduced feature that enables the translation of microscopy images into simulation templates significantly enhances the software’s usability and flexibility.

### Design and Implementation

#### Simulations in VirtualLeaf

Models in VirtualLeaf consist of several different layers that influence each other: the Cell-Based Model (CBM) for the *tissue*, the Individual-Based Model (IBM) for the behaviour of the individual *cells* and the Ordinary Differential Equation (ODE) models for intra- and intercellular signalling and biochemical processes, including intercellular diffusion of molecules. To keep a clear distinction between biological entities or processes and their computational equivalent, we will use *italics* to refer to the computational version. *Cells* in VirtualLeaf are represented as polygons composed of nodes connected by springs (see Section 1.3). *Tissues* are then formed by connecting *cells* into meshes. Adjacent *cells* share their nodes and springs, ensuring their relative position (the position of nodes relative to each other) cannot change except through specific moves. A Hamiltonian operator represents the *tissue*’s mechanical energy. This Hamiltonian comprises the compression of *cells* and the stretch/resistance of the *cell wall* and can be expanded to include other relevant elements, such as *cell* stiffness or a target *cell length*. Formulating the model in this way permits the definition of a hybrid model that operates across multiple time and space scales while coupling biochemical and mechanical dynamics (see Fig. 1).

**Fig 1.**
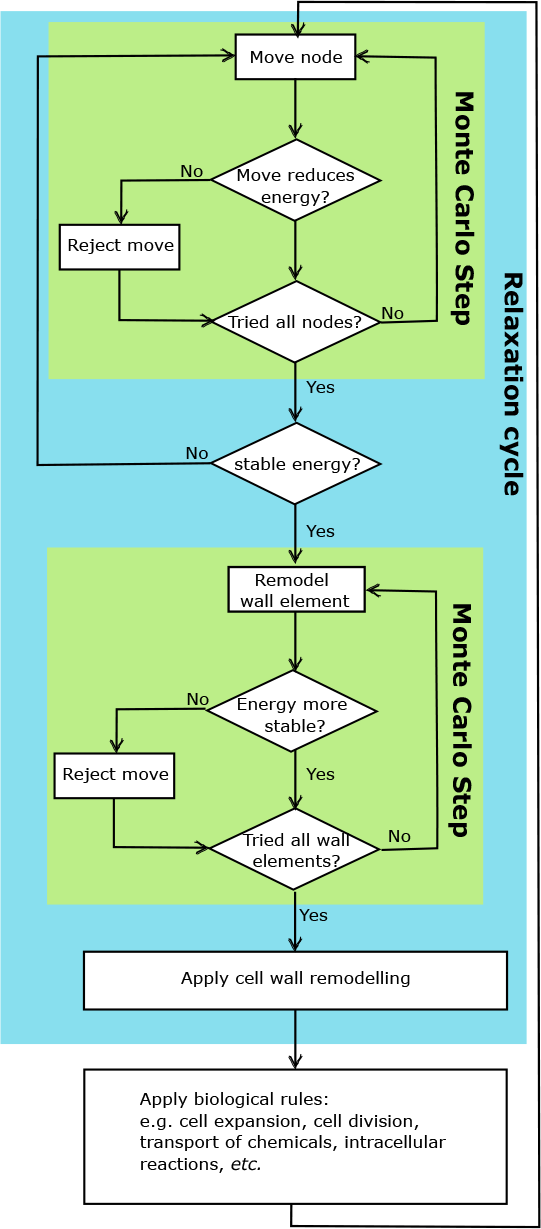
Updated overview of the simulation steps. After the Monte Carlo step for node displacement, there is a second Monte Carlo step assessing all *cell walls* per node w.r.t. *cell wall remodelling*.

For each simulation step, the energy change of the tissue is assessed using the Hamiltonian. The Hamiltonian (Δ*H*) is composed of a contribution of *cell* compression or area conservation (Δ*H*_*A*_) and *cell wall* stretching (Δ*H*_*L*_).

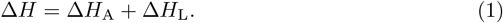

The area conservation contribution describes the *cell* ‘s turgor pressure and ensures the balance between cellular pressure and the pressure from the surrounding tissue [34]. It is defined as the sum of squared deviations of *cell area* (*A*(*i*)) from a resting area (*A*(*i*)_*t*_), weighted by a user-defined parameter (*λ*).

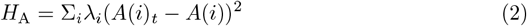

Similarly, *wall elements*, i.e. the *wall segment* between two nodes, are considered mechanical springs, with a resting length *L*(*j*)_*t*_ and a specific spring constant *λ*_*j*_:

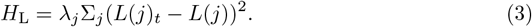

For a detailed introduction to the computational approach taken in VirtualLeaf and the installation instructions, we refer to [34] and [61]. The changes and new features that we introduce to VirtualLeaf in this paper concern the generation of *tissue* templates, the finer details of *cell wall* mechanics and the implementation of *cell wall* sliding.

##### 1.1 Tissue generation

An initial *tissue* layout must first be defined to simulate tissue dynamics and intercellular signalling processes. In VirtualLeaf, *tissues* are represented by a dynamic mesh of nodes that define *walls*, and non-overlapping polygons represent *cells* on the same mesh. VirtualLeaf uses an XML template that can be manually modified to simulate the desired starting configuration. Adapting the XML file can be a tedious process and, in this specific case, requires meticulous attention to ensure an accurate definition of *cell walls* over shared nodes. The updated version of VirtualLeaf was enhanced by incorporating a stand-alone, command-line SVG importer (*svg-to-vl* see Section 4.2) that can be used independently of VirtualLeaf. The importer facilitates the creation of the initial *cell* configuration through a user-friendly graphical interface, allowing users to visually model the *cells*’ spatial arrangement in the *tissue. Svg-to-vl* enables the streamlining of the domain definition phase, allowing the user to focus more on defining biological processes.

##### 1.2 Improved cell wall mechanics

Cell walls play an integral role in plant morphology and development: plant tissues are composed of cells of various ages and cell types that fulfil very different roles for plant survival. By defining *wall element*-specific stiffness *w*(*j*), we can now describe anisotropies in cell wall structures of a single cell, such as **cell-1** in Fig. 2a: The shaded top and bottom *cell wall elements* have different stiffness values than the vertical ones. During simulations, the specific stiffness for this *wall element* is integrated into the Hamiltonian; the base *cell*-specific value applies to the *wall elements* where no additional stiffness was defined Fig. 2b. We further refined the *cell wall* relaxation behaviour, calculating the Hamiltonian independent of the *target length* using a similar approach to [62]. Where [62, 63] rely on the resting length of the entire wall of a cell to assess the relaxation behaviour of the wall, we use the base length *B*(*j*) of every individual *wall element*, i.e. the length of the element at rest without any forces acting on it. In the Hamiltonian *H*_*L*_ itself, we use the degree of stretching to assess the energy in the system:

**Fig 2.**
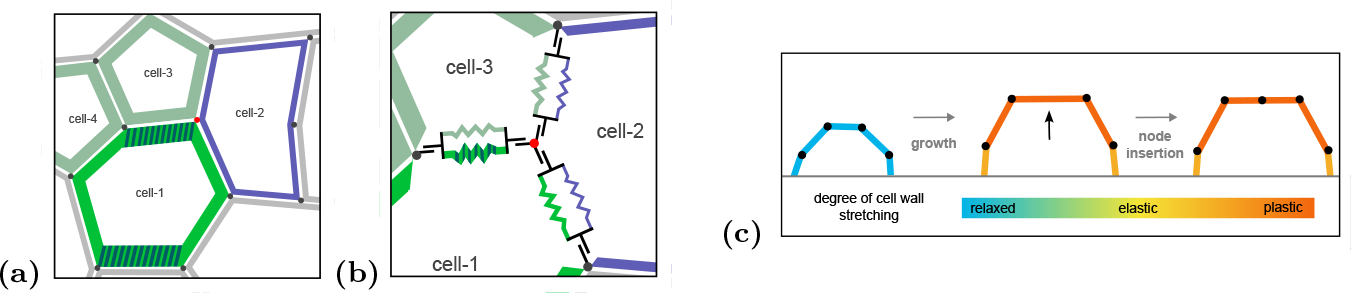
Refined *cell wall* mechanics in VirtualLeaf. (a). *Cells* can now have *cell*-specific *wall* stiffness values as indicated by the line thickness and colour of **cell-1** to **cell-4**. Additionally, *cells* can also have anisotropic wall stiffness values as indicated by the shaded wall elements of **cell-1**. (b) VirtualLeaf will take into account the stiffness values for all cell elements connected to a node when evaluating the change in the Hamiltonian during tissue simulations. (c) The introduction of new nodes no longer results in the relaxation of cell wall elements. The two new wall elements will remain under tension, maintaining the tissue layout [68].

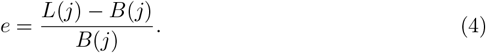

The cell wall component of the Hamiltonian is composed of *λ*_*L*_ (the general scaling factor for all cell walls), *E* (representing the elastic modulus of the tissue), the base length *B*(*j*), *w*(*j*) (the wall element-specific stiffness) and (*e*(*j*)_*t*_ − *e*(*j*))^2^ (the squared difference of the stretched positions of the cell wall element).

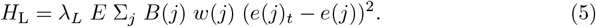

The fundamental principle of *tissue* simulation in VirtualLeaf remains unchanged. *Cell wall* elements are described as springs being stretched (or compressed) throughout simulations. The newly introduced aspect is that this calculation became independent of the target length and instead refers to the base length of the wall element. The deformation is assumed to be elastic for an extension of the wall element up to a certain threshold (*elastic limit*). The default value for this threshold is 15%, in the range of experimentally determined values (see Table 1). Beyond this threshold, experiments demonstrate cellulose fibres transition from elastic to plastic behaviour (i.e. they irreversibly deform). We represent this transition by updating the wall element’s base length if it extends beyond the threshold. Although we have removed the target distance between nodes (target length) from the cell wall’s contribution to the Hamiltonian, introducing new nodes still depends on the overall length of a wall element exceeding the defined threshold, a behaviour consistent with the previous releases of VirtualLeaf.

**Table 1.** Overview of elastic extension limit in different plant species.

| Species | Tissue | Elastic limit | Reference |
| --- | --- | --- | --- |
| *Zea mays* | | $\geq 18$ % | [64] |
| *Chara corallina* |  | 10 % | approximated from [65] |
| Sunflower |  | 12 % | [66] |
| Barley |  | 17.5 % | [66] |
| Dandelion |  | 6.9 % | [66] |
| *Aristolochia macrophylla* | outer layer | 2 % | [67] |
|  | whole axis | 3 % | [67] |
|  | inner chore | 3 % | [67] |

##### 1.3 Cell wall remodelling

While cell wall integrity and stability play an integral part in plant development and survival, there are very specific circumstances during a plant development where localised growth necessitates the extensive remodelling of the tissue, e.g. during lateral root emergence [54] and nodule development [69].

To implement this in VirtualLeaf, we must allow for an additional way of “node movement” during *tissue* simulations (Fig. 1). The nodes still move randomly to find a new, energetically more favourable position, as in previous versions of VirtualLeaf. To allow *wall elements* to “slide” to a more favourable node connection, we adopt a similar ‘sliding’ operator as previously proposed for modelling animal tissues [70]: In Fig. 3a, the depicted geometry results from the concurrent expansion of neighbouring *cells* along the outer border of the tissue domain and the discretisation caused by *wall elements*. By “sliding” *wall element* **b** outward by one node — thereby connecting it to **c** rather than **d** — the energetic tension in the simulation model is reduced, producing a more realistic transition.

**Fig 3.**
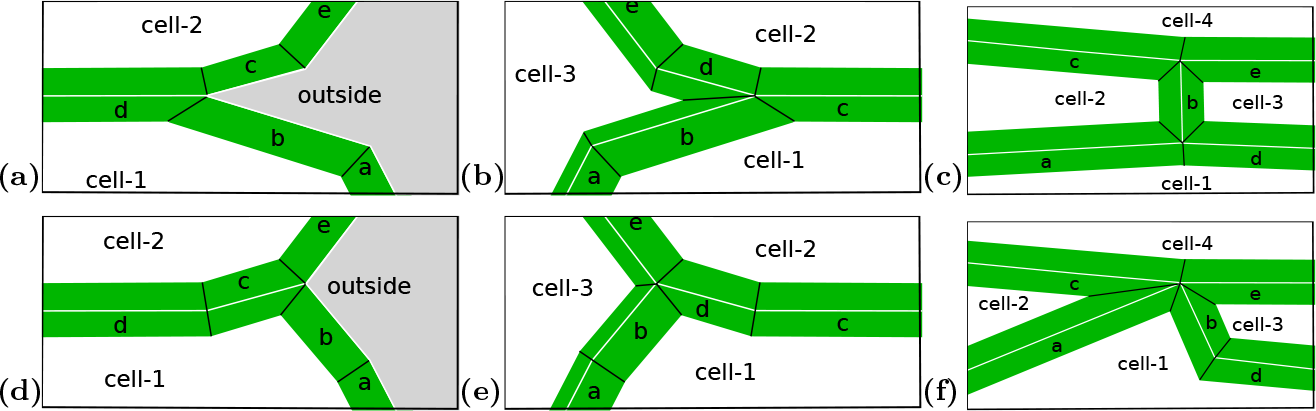
The representation of the behaviour of the *wall elements* between the *cells* in case of squished edges. These figures illustrate the system’s behaviour, whereas the decision-making process is described by Eq. (6). (a) During the growth of two neighbouring border *cells*, the closing of the space between **b** and **c** is imminent. (d) Moving the *wall element* **b** from the intersection between **d** and **c** to the intersection between **c** and **e** benefits the energetic equilibrium. (b) The **cell-3** is compressed between **cell-1** and **cell-2**, forcing the *cell wall elements* into a sharp angle (between **b** and **d**). (e) VirtualLeaf will move the *wall* between the lower and left cell (between **d** and **e**) to reduce the tension on the *cell wall*. (c) *Cell* compression on a *wall element* intersection will be detected by evaluating the *remodelling* Hamiltonian between **a** and **c** assuming the **b** *wall* removed. (f) The **cell-1** is exerting pressure on the thin and stretched **cell-2** and **cell-3**. The compression is relieved by the same algorithm as in (e) and (d), resulting in remodelling the wall-element **a** to the intersection of **b** and **c**.

After each *remodelling* event, the base length of each *wall element* is updated to prevent an artificially high strain on these *elements* (see Section 1.2). *Cell walls* are interconnected structures where changes in one region influence neighbouring areas due to shared mechanical forces. The *wall relaxation* (i.e. updating the base length) ensures the redistribution of forces, preventing imbalances and preserving the integrity of the model’s force network. This method reflects the current understanding of the dynamic nature of cell walls, aligning with plant cell behaviour under growth and mechanical strain [71]. In Fig. 3e the **cell-1** wall elements stiffness **b** and **d** are restored to their equilibrium.

A separate Monte Carlo step was introduced before *cell* housekeeping — the routines that update the simulation-specific *cell* status — and after the Monte Carlo node displacement to evaluate each wall element’s remodelling energy change. Unlike the approach taken by [70], however, this second Monte Carlo step is based on the *wall elements* and the Hamiltonian for *wall remodelling* is based on the spring energy in the *wall elements*. The *wall* between two *cells* on both sides of the *remodelled wall* changes in opposite directions, resulting in Eq. (6).

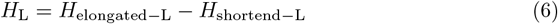

In cases where multiple potential *wall remodelling* events involve identical nodes (partial overlap), only one of these events will be permitted to proceed. This *remodelling* event is selected stochastically, based on the energy change involved. Since these *remodelling events* disrupt a *cell* ‘s list of *wall elements* and *neighbours*, the execution and selection of the *wall remodelling* events are addressed during *cell* housekeeping. In this simulation step, *cell division* and *growth processes* occur, which also impact the lists of *cell walls* and *neighbours*, necessitating a crucial repair step of these elements.

Furthermore, we have included the possibility for *cells* to “veto” *cell wall remodelling* events by deactivating *remodelling* events for specific *cell walls* or setting the *cell* ‘s property *veto remodelling*. Specific *cells* or *cell types* can be excluded from or included in *cell wall* remodelling. This property is accessible through the model in cell housekeeping and can change throughout the simulations. All *cells* sharing the remodelled *cell walls* can “veto” the *remodelling* events individually.

##### 1.4 Tissue simulations

VirtualLeaf (written in C++) was built on a local computer for the tissue simulations included in this manuscript. The model files and tissue XML files are included in the new release of VirtualLeaf; specifically, the highly simplified lateral root-like structure model employs the LateralRoot.xml template, and the severely minimized infection model utilises the pathogen infection.xml file. A pre-compiled version of VirtualLeaf, including the newly introduced models as well as additional models only visualising cell outlines and cell types, is provided as part of the new release for Linux, Windows and MacOS. The simulation results (both XML files and movies) are available as supporting information.

##### 1.5 Verifying Backwards Compatibility

The auxin growth model was simulated ten times in both the 2021 release [61] and the version presented here, utilising the new compatibility mode to test backwards compatibility (see S1 Video). The compatibility mode allows for the use of previous implementations of the Hamiltonian. As implemented in R, the resulting cell counts and area values were compared using the Kolmogorov-Smirnov and Wilcoxon rank sum tests (see Section 2.3).

## 2 Results

### 2.1 Initial Tissue Configuration (*svg-to-vl*)

With the new translation tool for microscopy images, we can now create realistic tissue templates for simulations in VirtualLeaf. Fig. 4 exemplifies this process for two different tissues, specifically the vascular cambium [72] and a longitudinal cut through the procambium [73]. Based on the microscopy images Figs. 4a and 4d, we created a traced image in Inkscape Figs. 4b and 4e, coding the different cell types by using different colours for the cell outline [74]. The created svg file is then used as input for *svg-to-vl* to create a tissue template for VirtualLeaf (see Fig. 4c and Fig. 4f). The tissue templates were used in subsequent test cases for the improved *cell wall mechanics* (see Section 1.2) and *cell wall remodelling* (see Section 1.3).

**Fig 4.**
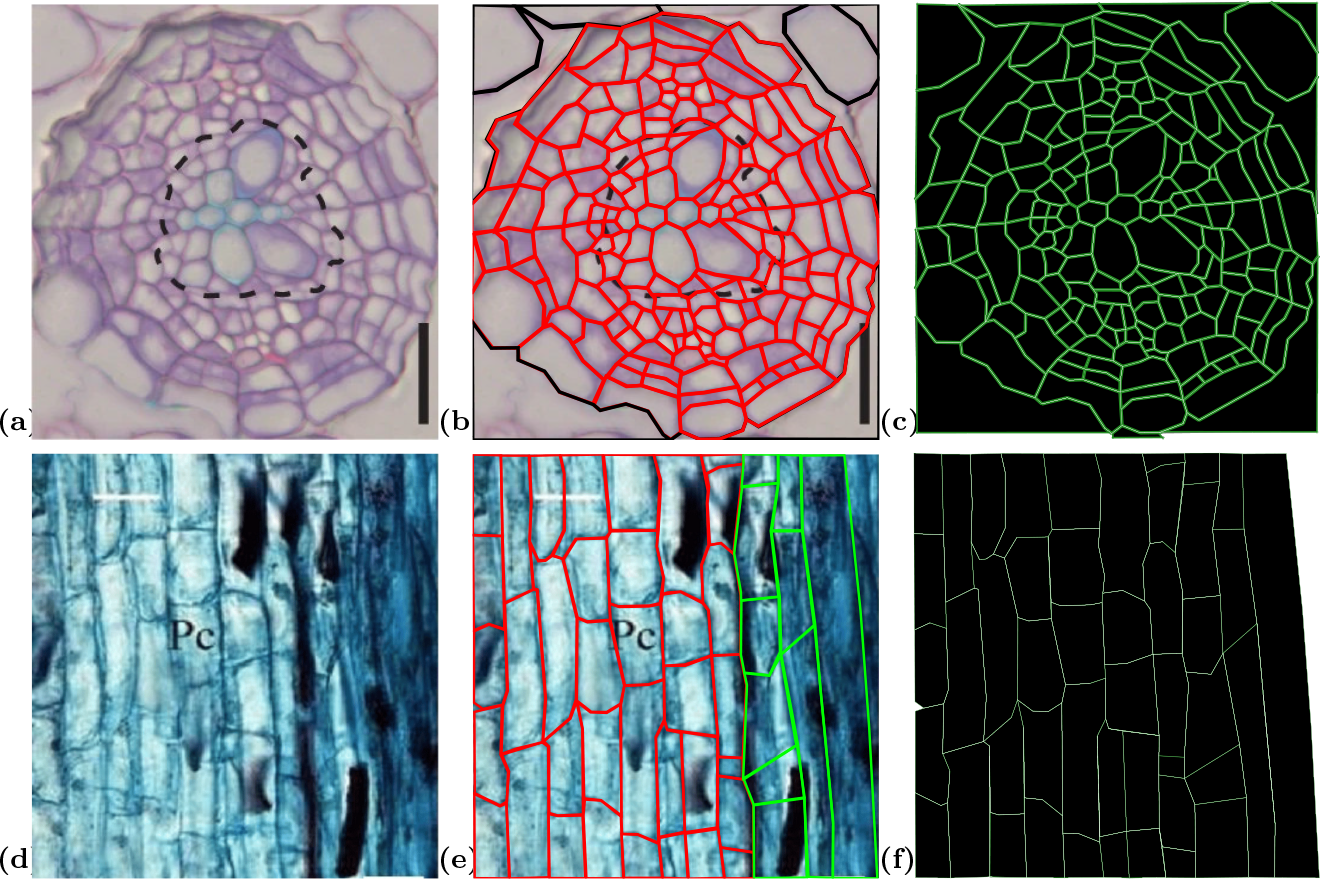
Representation of the process of tissue modelling in three stages, from microscope image to VirtualLeaf XML tissue data. (a) initial microscope image of plant tissue section from [72]. (b) The SVG drawing was produced with Inkscape [74], keeping Fig. 4a as the background image. (c) SVG drawing Fig. 4b converted to VirtualLeaf XML tissue. The black outlined cells are border cells fixed in place for the simulation. (d) initial microscope image of longitudinal plant tissue (procambium) from [73]. (e) The SVG drawing was produced with Inkscape [74], keeping Fig. 4d as the background image. Xylem cells (Green) are differentiated from procambium cells (red), giving structural integrity to the vessel wall. (f) SVG drawing Fig. 4e converted to VirtualLeaf XML tissue.

### 2.2 Improved cell wall mechanics

The refined *cell wall* mechanics of VirtualLeaf open up several new research directions where a more detailed mechanical description is necessary, e.g. during pathogen infection. Pathogens use various strategies to weaken the plant cell wall during infection, such as the secretion of cell wall degrading enzymes by plant-pathogenic fungi [75, 76, 77, 78]. In turn, pathogen detection by the plant triggers a wave of Ca^2+^ and reactive oxygen species (ROS) along the plant tissue [79], resulting in a more stable interaction between cell wall filaments [80]. The presented version of VirtualLeaf is able to simulate these cell wall infection dynamics. To illustrate this, we built a simplified model of infection dynamics in a small tissue region (see Figs. 5a to 5d and S3 Video), which was generated using *svg-to-vl* (see Fig. 4f). Here, a foreign cell (e.g. a fungi) starts growing at the outside of the tissue. Through the release of *cell wall* destabilising components, the *cell wall* s are slowly weakened (as illustrated by the decreasing line thickness) until the *pathogen* can infiltrate the *plant tissue*.

**Fig 5.**
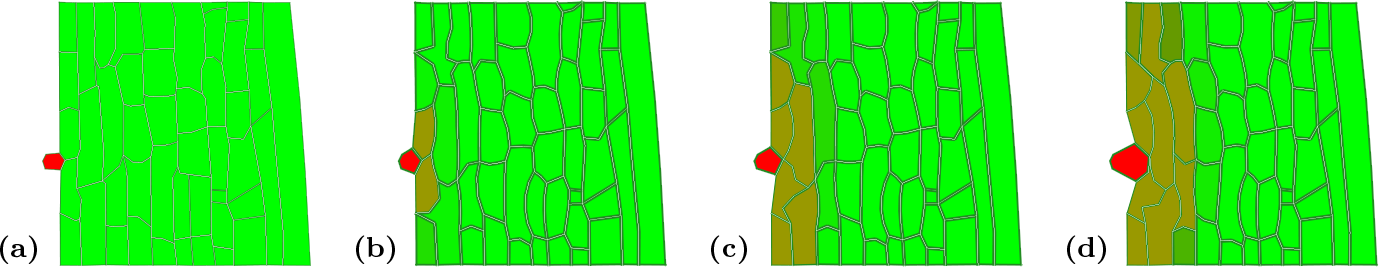
Time-lapse visualisation depicting a pathogen compromising tissue integrity. The pathogen initially sits at the outside of the tissue (a). As the pathogen grows, it exudes a cell wall weakening chemical ((b) and (c), shown in brown), weakening the cell walls and destabilising the tissue. This allows the pathogen to infiltrate the plant tissue (d).

### 2.3 New implementation of cell wall dynamics is compatible with older models

While we sought to improve cell wall mechanics in VirtualLeaf, we also wanted to ensure the compatibility of this updated version of VirtualLeaf with models built using previous versions of VirtualLeaf. We set all default values for the parameters related to cell wall stability so that the tissue dynamics simulations of existing models remain unchanged. We verified this behaviour by comparing the simulation results between VirtualLeaf v2.1.0 and v1.1.0. The simulations were repeated ten times for both software versions. The respective cell counts and area values were compared statistically with a visual inspection of the resulting cell tissue. Neither cell count nor cell area showed a significant difference between simulations based on a Kolmogorov-Smirnov test (cell count: p-value = 0.3663; cell area: p-value = 0.4175) and the Wilcoxon rank sum test (cell count: p-value = 0.1728; cell area: p-value = 0.1431).

### 2.4 Cell wall remodelling

While cell wall stability is fundamental to plant stability and survival, there are instances during plant development in which the cell wall is actively remodelled (see Section 1.3). The example model illustrates this process within a simplified framework (see Fig. 6). In the simulation, a single cell is designated as the founder cell initiating the lateral root-like structure. For the sake of simplicity, only this cell undergoes growth, and upon division, the daughter cell furthest to the right assumes the role of the new initiator. This minimal condition suffices to generate a structure that loosely resembles a lateral root.

**Fig 6.**
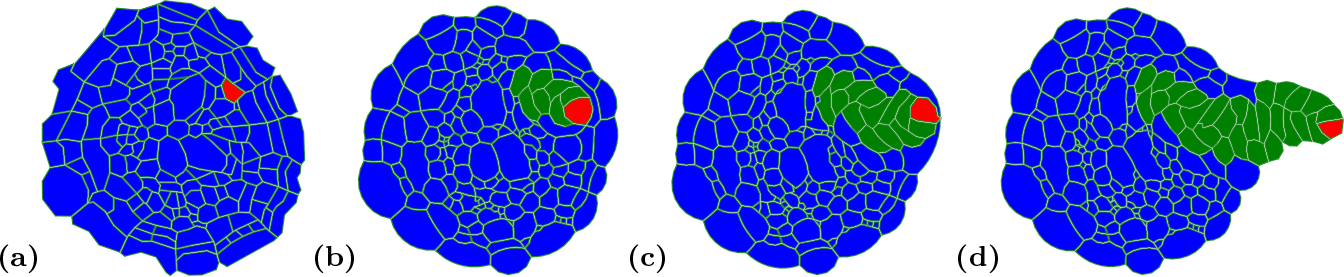
Time-lapse visualisation of a proliferating lateral root-like structure emerging from the stationary (assumed non-growing) taproot, as depicted in Fig. 4a. The cell colour distinguishes cell types. Dynamic remodelling of wall elements facilitates the migration and expansion of growing cells through the quiescent cell population within the taproot tissue. The underlying simplified model includes only one cell undergoing growth and division. In each division, the rightmost cell is selected to grow next. The time-lapse sequence is extracted from the video in the supporting information (S2 Video). (a) Initial simulation state of the quiescent tissue (refer to Fig. 4c). A single cell in the fourth cell layer is designated as the origin of the lateral root. (b) As the growing cell cluster expands, it exerts pressure on the surrounding quiescent tissue. (c) The expanding cell cluster reaches the boundary of the tissue. (d) Remodelling of wall elements facilitates the growth of the lateral root-like structure through the stem wall, allowing it to emerge outside the taproot.

Dynamic wall-element remodelling allows cell walls to adjust by shifting one wall-element ending between nodes, evading pressure from neighbouring cells. By facilitating these adjustments, cell wall remodelling prevents the formation of cell shapes that are energetically unfavourable and unlikely to occur naturally. Wall-element remodelling allows cells to squeeze between other cells if energetically profitable, for example, supported by the pressure of a growing tissue cluster (see Fig. 6). The video in the supplementary section S2 Video shows a more derailed evolution of the time-lapse represented in Fig. 6.

This adjustment is particularly beneficial in scenarios where different spatial cell clusters exhibit varying growth rates, such as during callus formation in response to wound healing [81], in root nodules containing nitrogen-fixing bacteria [69], and in gall formation initiated by insect activity [82].

## 3 Availability and Future Directions

VirtualLeaf fulfils a vital role in facilitating the simulation of plant tissues [6, 60], particularly in areas currently inaccessible to life-cell imaging where computational modelling can provide an indication for describing the dynamics of developmental processes such as radial growth [6]. Furthermore, it has been the starting point of several new computational approaches [70, 83]. While our updates to VirtualLeaf represent an essential step towards accurately simulating plant tissue growth, some limitations still need to be considered. Important effects such as growth anisotropy can currently only be represented by prescribing growth direction [60]. VirtualLeaf is currently actively developed further in our groups and those of others. It is expanded to new research areas, such as plant mechanobiology, brassinosteroid signalling, and nodulation. Importantly, VirtualLeaf is currently limited to two-dimensional simulations, whereas for many problems in plant development, the three-dimensional context of cells is crucial. We are, therefore, currently extending VirtualLeaf to three dimensions and implementing additional biomechanical components to support anisotropic growth. All in all, these developments will give VirtualLeaf capabilities to simulate a more comprehensive array of biological processes to support hypothesis-driven research in plant biology.

## Supporting information

### Video and Image generation

VirtualLeaf generates PDF and PNG images at regular intervals during the simulation (named *leaf* [0 − 9]^+^.[*pdf* | *png*]). These images were converted using the “convert” image conversion package available on Linux. Subsequently, the sequence of images was compiled into videos using the FFmpeg software. The following script was employed to automatically convert a directory of VirtualLeaf images into videos and their corresponding EPS file versions.

#### Listing 1.

bash 5.1

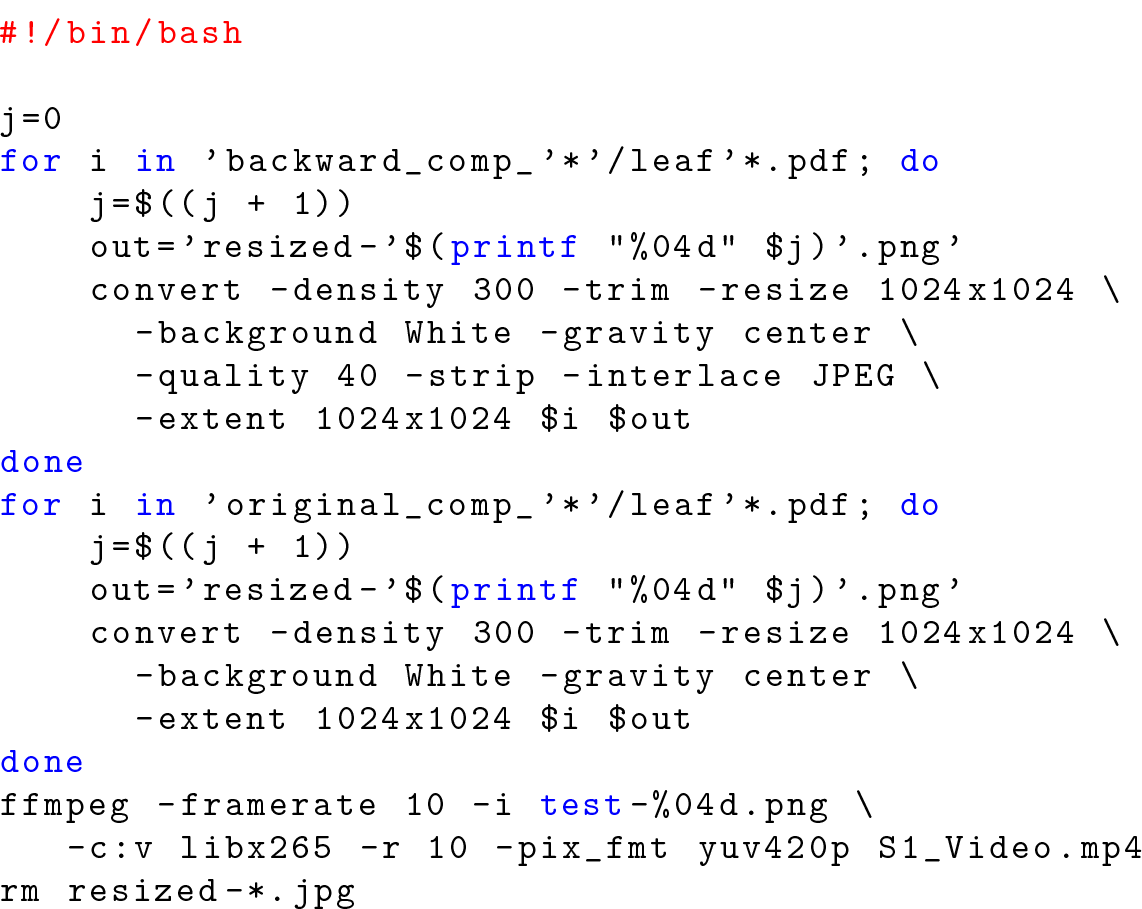

**S1 Video.**
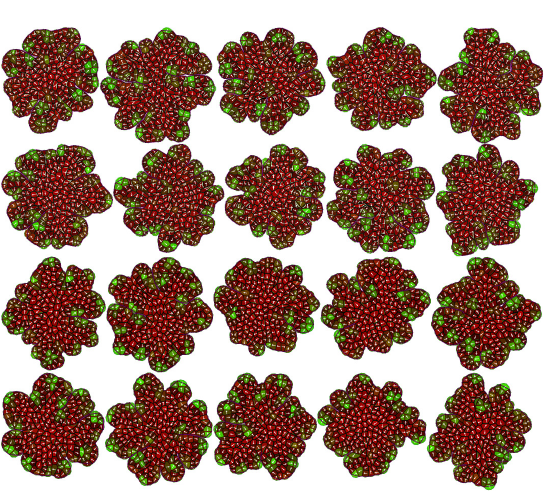

#### 10 times random growth of the auxin model updated version followed 10 times random growth of the auxin model in the previous version

The S1 Video, titled “S1 Video.mp4,” is included in the supplementary materials. This video was generated using data from the “virtualleaf simulation data v2.1.0.zip” file, specifically within the “compatibility” directory. The results from the two sets of ten compatibility simulation runs are organised into separate subdirectories. The S1 video is a sequential compilation of all images from these simulations, providing a visual summary of the entire dataset.

**S2 Video.**
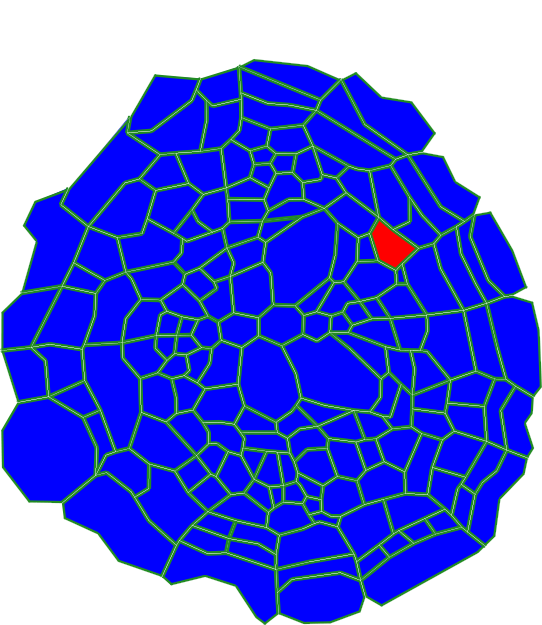

#### Growth dynamics during lateral root-like structure initiation including cell wall loosening

The S2 Video, titled “S2 Video.mp4,” is included in the supplementary materials. This video was generated using data from the “virtualleaf simulation data v2.1.0.zip” file, specifically within the “lateral root” directory. The simulation has been based on the “import/root draw plain.xml” file. Available in the binary distribution as “data/leaves/lateralRoot.xml” as default for the Model “Lateral Root Growth”. The subsequently generated images were concatenated to the “S2 Video.mp4” file.

**S3 Video.**
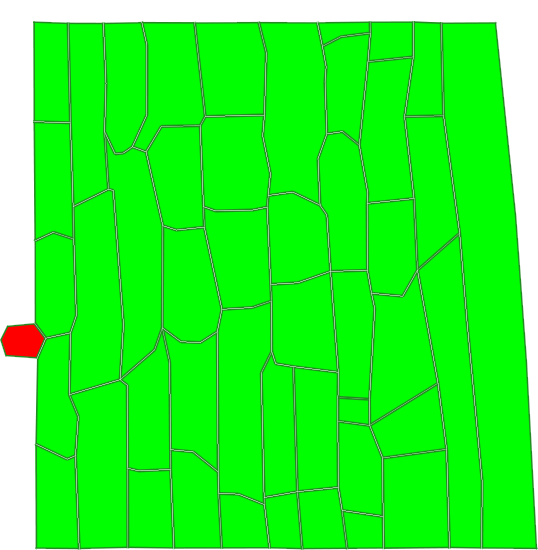

#### Pathogen growth and cell wall weakening effects in VirtualLeaf

The S3 Video, titled “S3 Video.mp4,” is included in the supplementary materials. This video was generated using data from the “virtualleaf simulation data v2.1.0.zip” file, specifically within the “infection growth” directory. The simulation has been started from the “imports/long draw plain.xml” file. Available in the binary distribution as “data/leaves/pathogen infection.xml” as default for the Model “Infection”. The subsequently generated images were concatenated to the “S3 Video.mp4” file.

## 4 *svg-to-vl* Instructions

This updated release of VirtualLeaf introduces a command-line tool (*svg-to-vl*) designed to simplify the conversion of plant tissue models into simulation-ready templates, referred to as “leaves.” These highly adaptable templates extend beyond leaf tissues to various plant structures, enabling broader applications in plant modelling. The release also includes a comprehensive, step-by-step guide for using the tool, including creating and formatting scalable vector graphics (SVG) files required by the script.

### 4.1 Prerequisites

For the script to function properly, ensure Python [84], pip and Inkscape [74] are installed on the personal computer (Listing 2.

#### Listing 2.

Open a terminal or command prompt and run:

~~~
python -- version
pip -- version
~~~

These commands verify their availability and versions. The minimal tested versions for compatibility with *svg-to-vl* are Python 3.10 and pip 22.0.2. Use the pip install command to install *svg-to-vl* (Listing 3).

#### Listing 3.

Open a terminal or command prompt and run:

~~~
pip install svg - to - vl
~~~

This command will download and install the program and its dependencies (shapely [85], numpy [86], simpy [87]) from the Python Package Index (PyPI).

### 4.2 Generating the SVG File

In this step, a microscopy image serves as the foundation for creating an SVG representation of the tissue, which is subsequently converted into an XML simulation template (referred to as a “leaf”) using *svg-to-vl*. The process involves drawing freehand paths in Inkscape to delineate the circumference of each cell. These boundaries provide coordinate points that uniquely define each cell’s geometry. *svg-to-vl* translates these coordinates into nodes representing each cell in the template.

Individual cell types can be assigned by applying distinct colours to the paths during this stage. For instance, epidermal cells may be coloured red, while xylem cells can be designated blue. Fixed cells are predefined with a black outline in the script, which can be modified if necessary. For simplicity and consistency, we recommend using black to outline the fixed cells of the tissue. This approach clarifies the definition of structural and functional roles within tissue simulation.

The cells in the SVG image must be traced using a closed path to generate the tissue template provided in the supplementary materials (Fig. 7). Begin by selecting the pen tool (designed for creating curved and straight lines) in Inkscape, then mark critical points along the cell boundaries. These points will later be converted into node coordinates, so it is essential to capture all significant details of each cell’s geometry accurately.

**Fig 7.**
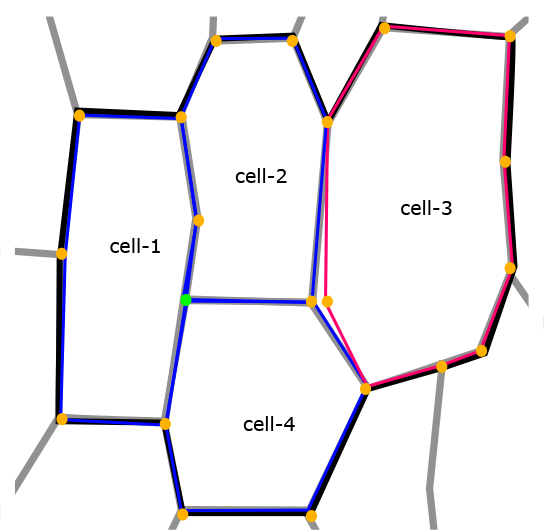
Generating an SVG template file for the Python script requires careful tracing of the tissue outline. Several constellations can cause issues in the translation from SVG to XML: While tracing the microscopy image, special care should be taken to include all essential points. While the highlighted green node might not be required for the outline of **cell-1**, it is necessary to define the boundary between cell-2 and **cell-4** and should, therefore, also be included in the outline of **cell-1**. Furthermore, the corner points must be close together for neighbouring cells to be recognised as one node. Otherwise, a double node situation occurs, as illustrated in **cell-3**.

Ensure that each path starts and ends at the same point to form a closed shape; this is critical for the software to recognise cells during the subsequent processing. Assign a uniform colour to cells of the same type while tracing to predefine their classification at this stage. For example, all cells of a specific type, such as epidermal cells, should share the same colour to streamline later identification and simulation tasks. This systematic approach ensures precise cell type definition and geometry for the simulation template.

In the example tissue, **cell-1, cell-2**, and **cell-4** have the same cell type and are therefore outlined in the same colour (blue). **Cell-3** belongs to a different cell type and is therefore outlined in red. During the tracing process, the essential points for that cell but also those of the neighbouring cells need to be considered: While the green node in Fig. 7 is not necessary for **cell-1**, it is essential for defining the border between **cell-2** and **cell-4** and must be included in the outline of all three cells. The shared points between cells must be placed at identical spots or at least very close together to be recognised as one node. Otherwise, a situation will occur, such as with **cell-3** in Fig. 7, where there is a gap in the tissue. This erroneous gap will result in a small erroneous cell being defined in the translation process.

The two generated example XMLs and the appropriate SVG files used for S2 Video and S3 Video videos are available in the supplementary data file “virtualleaf simulation data v2.1.0.zip”, specifically within the “imports” directory.

The file “long draw plain.xml” was generated from “long draw plain.svg”. The file “pathogen infection.xml” was generated from “root draw plain with ptho.svg”.

### 4.3 Running *svg-to-vl*

*svg-to-vl* facilitates the customisation of multiple arguments, including the designation and directory of the template file. Users can specify a scaling factor to map x and y coordinates to the cellular scale and a colour map for encoding cell types within the SVG file. Specific details for each cell type are provided as arguments, delineated by colons and their hex RGB colour code, encompassing elements such as cell type and intracellular chemical concentrations.

*svg-to-vl* -i “path to the SVG file (without .svg)” -t “path to the XML template file (with file extension)” -s “numerical scaling factor between image and simulation template” -c “colour code”

The colour code has the form of a hexadecimal RGB colour code, cell type, intracellular chemical concentrations: “ffffff,1,2.25,0.48:ff0000,3,2.5,1.5” until all colour codes are described.

#### Listing 4.

Complete example command

~~~
svg - to - vl -i “imagefile “-t “template. xml” \
    -c ffffff, 1, 2.25, 0.48: ff0000, 3, 2.5, 1.5
~~~

## 5 Control of Features

To ensure seamless compatibility with prior versions of VirtualLeaf, we have implemented an augmented parameter named “compatibility level” within the option framework. This parameter operates as a bit flag; additional features are (de)activated by (re)setting the bit value in the “compatibility level” unsigned 16-bit integer value. This “compatibility level” also facilitates the seamless integration of forthcoming enhancements. The “WALL STIFFNESS HAMILTONIAN” constant is represented by the value “1” (corresponding to the bit value “0×0001”), while the “WALL SLIDING” corresponds to “3” (bit value “0×0003”. The wall sliding functionality necessitates the activation of wall stiffness. The default value is presently set at “65535” (corresponding to a bit value of “0xFFFF”), thereby ensuring the activation of supplementary future features.

## 6 Example Simulations

### 6.1 Infection model

For this highly simplified infection model, the tissue template shown in Fig. 4f is used as the foundational framework, deliberately omitting many biological complexities for conceptual clarity. A pathogenic cell (e.g., a fungal cell) was added to the outside of the tissue. The outside boundaries of this tissue are fixed to maintain the overall tissue shape, and all remaining nodes can move according to the tissue energy. The pathogenic cell secretes a chemical factor that diffuses throughout the tissue and degrades cell wall stability, while plant cells degrade this chemical at a rate of *k*. The pathogenic cell will grow and use the destabilised cell walls to grow into the plant tissue.

The intracellular model only considers the dynamics of the pathogenic chemicals. In the pathogen, we assume a constant chemical level for simplicity’s sake. For plant cells, the actual chemical levels will depend on the diffusion and the degradation within the cell:

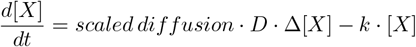

#### 6.1.1 Scaling factor for pathogenic chemical diffusion

In the equations describing the pathogen diffusion from one cell to the next, we have included a scaling factor for the pathogen concentration changes in **cell-1** and **cell-2**. To ensure mass conservation, we require that the total amount of pathogen remains constant:

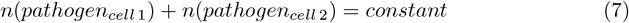

and as a consequence

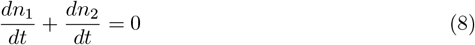

It follows that

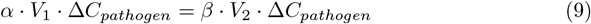

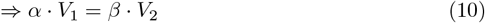

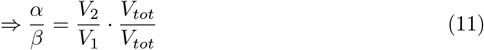

with *V*_*tot*_ being the sum of *V*_1_ and *V*_2_. Based on the equations above, we set the scaling factors to

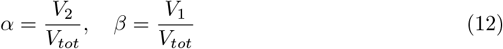

### 6.2 Lateral Root Model

The lateral root-like structure model, designed explicitly for conceptual clarity, begins with the imported taproot model (see Fig. 4a). In this intentionally oversimplified framework, a single cell is designated as the lateral root founder cell (cell type = 3), significantly reducing the complexity of biological processes. In this model, only the rightmost cell of cell type 3 is defined as actively growing. The cell division threshold is set to two times the average model cell area. During cell division, the rightmost daughter cell is selected to continue as the growing cell.

## 7 Source Code

VirtualLeaf is freely available as open-source software at GitHub under https://github.com/rmerks/VirtualLeaf2021. This manuscript is based on release v2.1.0 and is also available as a precompiled package at Github Releases (https://github.com/rmerks/VirtualLeaf2021/releases/tag/v2.1.0).

## 9 Author contributions statement

Ruth Großeholz and Richard W. van Nieuwenhoven contributed equally to the design and implementation of the research, the analysis of the results and the writing of the manuscript. Bruno Hay Mele contributed to the design of the cell-specific cell wall stiffness and reviewed the manuscript. Roeland M.H. Merks helped with the design in general and edited the manuscript.

## 10 Acknowledgements

R.G. would like to thank the CRC-1101 project D02 of the German Research Foundation and the Dutch Sectorplan 2 beta for funding and the Joachim Herz Foundation for its support. The authors acknowledge TU Wien Bibliothek for financial support through its Open Access Funding Programme. R.W.N. would like to acknowledge Ille C. Gebeshuber for her guidance, mentorship, academic insights and support, which have been invaluable throughout this research project. R.W.N. extends his gratitude to Fabian Buranits for his assistance in identifying Hamiltonian and programming problems during this research; his contribution and dedication to the project have been greatly appreciated. R.M.H.M. was supported by NWO grant NWO/ENW-VICI 865.17.004 and by the Dutch Research Council (NWO) through the Gravitation program GreenTE (project no. 024.006.001).

